# Aberrant synaptic release underlies sleep/wake transition deficits in a mouse *Vamp2* mutant

**DOI:** 10.1101/2020.01.09.900233

**Authors:** Gareth T. Banks, Mathilde C. C. Guillaumin, Ines Heise, Petrina Lau, Minghui Yin, Nora Bourbia, Carlos Aguilar, Michael R. Bowl, Chris Esapa, Laurence A. Brown, Sibah Hasan, Erica Tagliatti, Elizabeth Nicholson, Rasneer Sonia Bains, Sara Wells, Vladyslav V. Vyazovskiy, Kirill Volynski, Stuart N. Peirson, Patrick M. Nolan

## Abstract

Sleep-wake transitions are modulated through extensive subcortical networks although the precise roles of their individual components remain elusive. Using forward genetics and *in vivo* electrophysiology, we identified a recessive mouse mutant line characterised by a reduced propensity to transition between all sleep states while a profound loss in total REM sleep time was evident. The causative mutation, an Ile102Asn substitution in VAMP2, was associated with substantial synaptic changes while *in vitro* electrophysiological investigations with fluorescence imaging revealed a diminished probability of vesicular release in mutants. We conclude that the synaptic efficiency of the entire subcortical brain network determines the likelihood that an animal transitions from one vigilance state to the next.

---

Despite advances in our understanding of the neurophysiology of sleep (*1, 2*), the genetic regulation of its fundamental vigilance states – wakefulness, non-REM (NREM) sleep and REM sleep – remains poorly understood. An extensive subcortical circuitry of neuromodulatory nuclei is thought to regulate global sleep-wake transitions, but the specific role of individual components remains to be determined. While high-throughput forward genetics screening has provided invaluable insights into the molecular genetic mechanisms underlying circadian rhythms (*3-5*), traditional electroencephalographic (EEG) methods of studying sleep are not conducive to high-throughput approaches and currently only a single study using such an approach has been published (*6*). Here, we adopted a high-throughput hierarchical approach, initially using behaviourally-defined sleep prior to EEG/EMG to identify mutant pedigrees with abnormal sleep-wake parameters in *N*-ethyl-*N*-nitrosourea (ENU) G3 pedigrees. Cloning and sequencing of the strongest phenodeviant pedigree identified a mis-sense mutation in the transmembrane domain of VAMP2, the core vSNARE protein mediating synaptic vesicle fusion and neurotransmitter release. EEG analysis confirmed the reduced sleep phenotype and revealed a marked decrease in REM sleep. Furthermore, while the EEG signatures of the different vigilance states were largely unaffected, VAMP2 mutant animals showed a profound deficit in their capacity to switch states once a specific state had been initiated. To determine how such a previously unexplored phenotype may arise we show, using cellular, molecular, imaging and electrophysiological studies, that vesicular release efficiency and short-term plasticity are drastically affected in mutants. The consequences of this deficit in neuronal firing and the inherent inertia in state switching demonstrates a hitherto uncharacterised role for VAMP2 in sleep and highlights how new aspects of gene function, even for well-characterised genes, continue to be uncovered using forward genetics.

To identify mouse lines with sleep deficits, we used home-cage video-tracking to measure immobility-defined sleep (*7*) in cohorts from G3 pedigrees generated in a large ENU mutagenesis programme (*8*). By plotting percentage time spent immobile during light and dark phases (**Fig. 1A,B**) we identified a pedigree, called restless (*rlss*, MGI:5792085), where multiple individuals expressed reduced immobility. Differences were particularly evident during the light phase and were confirmed by screening a second cohort from the same pedigree. Further analysis in 1 hr time bins indicated a strong effect throughout the first eight hours of the light phase and towards the end of the dark phase (**Fig. 1C**). Using DNA from affected individuals, we mapped the non-recombinant mutant locus to a 35 Mb region on Chromosome 11 (**Fig. 1D**). Whole genome sequencing revealed a single high-confidence coding sequence mutation within the non-recombinant region. Sequence validation in multiple affected individuals consistently identified a single coding sequence variant co-segregating with the mutant phenotype, a T441A transversion in *Vamp2 (Syb2)* resulting in an Ile102Asn substitution in the protein’s transmembrane domain (**Fig. 1E,F**). VAMP2 is the major neuronal vesicular component of the Soluble *N*-Ethylmaleimide-sensitive Factor (NSF) Attachment Protein Receptor (SNARE) complex, fundamental to neurotransmitter exocytosis. Outcrossing affected individuals to wild-type mates confirmed the recessive phenotype, with immobility-defined sleep in heterozygotes being no different to wildtypes. Statistical analysis of immobility in 1hr bins confirmed a significant genotype effect and pairwise comparisons confirmed that sleep in homozygotes was significantly reduced in the first seven hours of the light phase and the final seven hours of the dark phase (**Fig. 1G**; for this and all other statistical analysis, refer to **Table S1**). Further analysis of mutants also identified unusual sleep behaviours where individuals showed no preference for nesting in sheltered parts of the home-cage, frequently moved sleep position within the cage and assumed atypical sleep postures (**Fig. 1H**). Conventional circadian parameters using cages with wheels could not be reliably measured in homozygotes (**Fig. 1J**). However, Passive Infrared monitoring of circadian rhythms (*9*), demonstrated that homozygote circadian period was no different from controls while period amplitude and activity parameters were affected (**Fig. 1J,K**).

**Figure 1.**
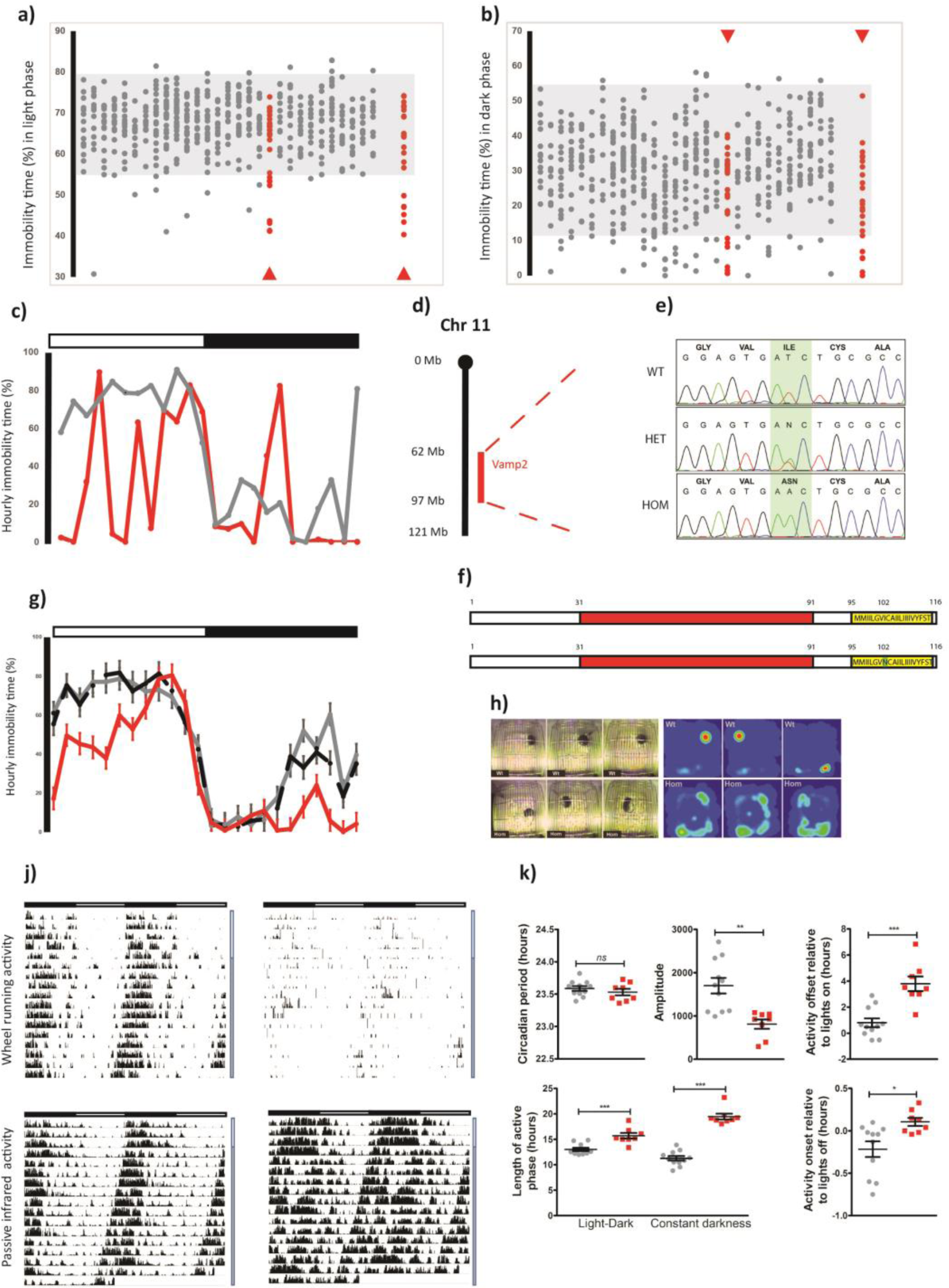
ENU sleep duration screen identifies a G3 pedigree with a mutation in *Vamp2.* Percent immobility-defined sleep during light **a)** and dark **b)** phases. Individuals from G3 pedigrees are shown in columns. Shaded areas represent normal immobility range. Arrowheads indicate the first and second cohorts of the pedigree. **c)** Percentage hourly immobilities for an affected individual (red) and unaffected littermate (black). Light (open bar) and dark (filled bar) phases are indicated above the graph. **d)** Mapping of affected individuals identified a 35 Mb region on Mmu11 including the *Vamp2* locus. **e)** Whole genome sequencing identified a single mutation in affected animals resulting in an lle102Asn substitution in the transmembrane region (yellow) of VAMP2 **f)**, SNARE motif shown in red. **g)** Hourly immobility percentages in homozygotes (red), heterozygotes (black dashed) and wildtypes(grey). **h)** Sleep patterns from video stills and heat maps over the duration of the light phase in wildtypes(top panels) and homozygotes (bottom panels). **j)** Double-plotted actograms of wildtypes (left panels) and homozygotes (right panels). Top panels show wheel-running activity, bottom panels show movement using passive infrared (PIR) sensors. Light and dark horizontal bars represent periods of light and dark where appropriate. Vertical bars to the right represent periods of 12:12 light dark (white)or constant darkness (grey). **k)** Circadian measures in wildtypes (grey) and homozygotes (red) calculated from PIR data. Individual data points are shown as is mean ± SEM, p<0.05*, p<0.01**, p<0.01***.

To investigate sleep-wake architecture in *Vamp2*^*rlss*^ mice, we implanted cortical EEG and nuchal EMG electrodes (**Fig. 2A**) and performed 24-h baseline home cage sleep recordings (**Fig. 2B**). EEG/EMG signatures were typical of wakefulness, NREM and REM sleep (*10*), although more pronounced slow-wave activity (0.5-4Hz) during NREM sleep and slower theta oscillations during REM sleep were noted in *Vamp2*^*rlss*^ (**Fig. 2C, Fig. S1A**). However, over the entire 24-h period *Vamp2*^*rlss*^ mice accumulated 1.4 hours (14%) less NREM sleep and 1.2 hours (57%) less REM sleep, with a ratio of NREM sleep time per wake time significantly reduced (**Fig. 2D**). Plotting the distribution of vigilance states across 24-h showed the difference between genotypes was especially apparent for REM sleep (**Fig. S1B, S2A**), a conclusion supported by a decreased REM-to-total sleep ratio (**Fig. 2E**), while the decrease in NREM sleep during the dark period was partially compensated during the second half of the light phase (**Fig. S1B, S2A**). Investigations as to whether reductions in sleep are related to a reduced probability of state transitions showed that the number of wake-to-REM transitions was markedly reduced in homozygotes (**Fig. 2F**). Moreover, increased wakefulness and longer wake episodes in homozygotes (**Fig. 2D, 2G, Fig. S2B**) suggested increased wake state continuity. In considering all wake episodes other than brief awakenings (≤16 sec), their frequency was halved in *Vamp2*^*rlss*^ mice, suggesting that once wakefulness is initiated it is sustained for longer durations. Similar dynamics also manifested in state transitions within sleep; once a NREM sleep episode was initiated, it was less likely to transition into REM sleep (**Fig. 2H, Fig S2C**). Almost 90% of all NREM episodes terminated in REM sleep in WT mice, compared to less than 70% in *Vamp2*^*rlss*^ mice (**Fig. 2J**). This indicates that, along the continuum of wake -> NREM -> REM occurrences, the inertia to transition to the next state is increased in *Vamp2*^*rlss*^ mice (**Fig. 2K**). Consistently the distribution of episode durations for NREM and REM sleep showed an increased incidence of longer episodes for both sleep states in *Vamp2*^*rlss*^ mice (**Fig. 2L,M**), further suggesting that once a specific state is initiated it is less likely to terminate.

**Figure 2.**
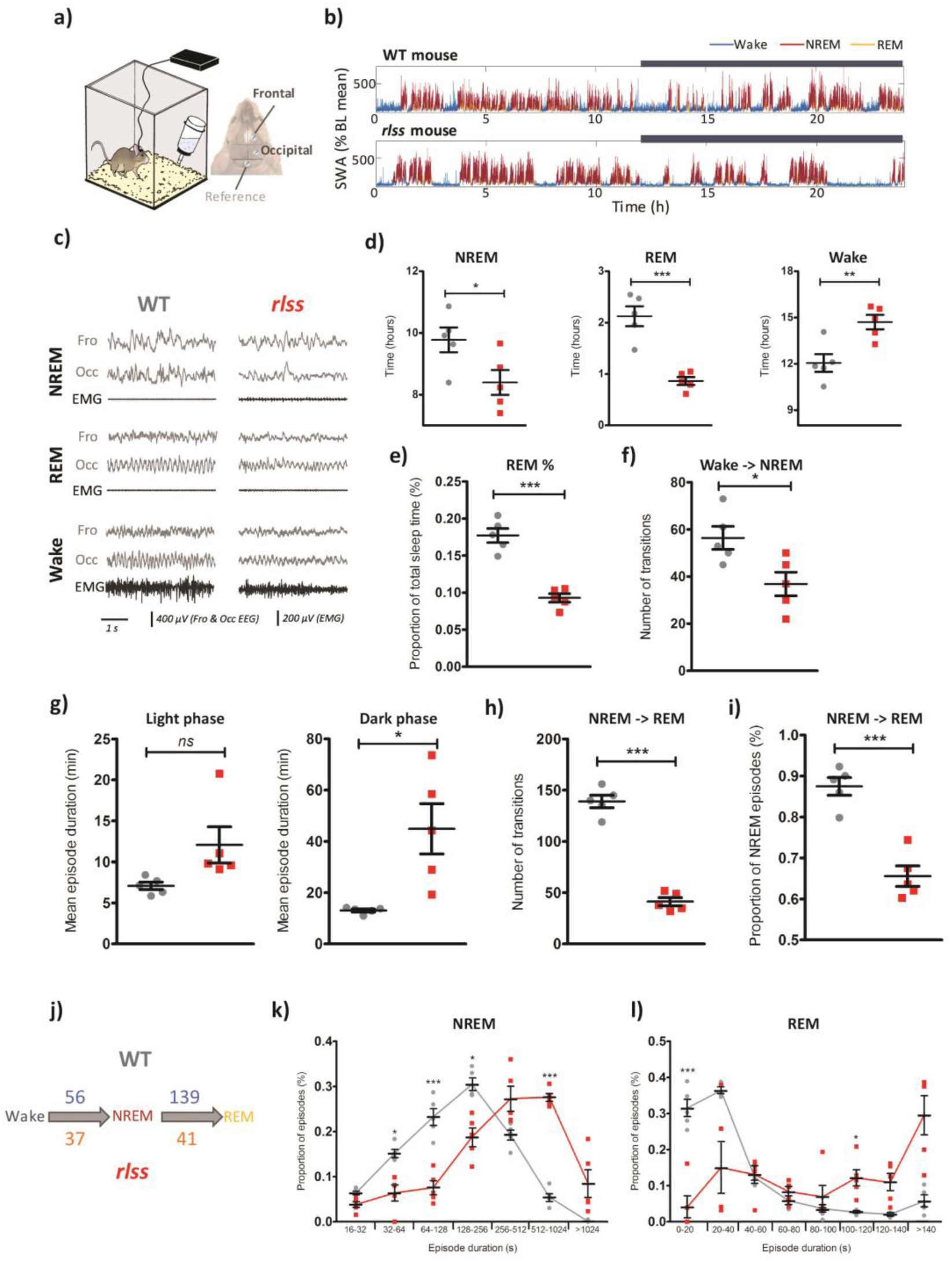
EEG abnormalities in *Vamp2*^*rlss*^. **a)** A tethered EEG set-up was used to perform chronic recordings from frontal and occipital cortex. **b)** Slow-wave activity (SWA) levels across 24h in individual WT and *Vamp2*^*rlss*^ mice. SWA levels are expressed as percentage of the 24h baseline (BL) mean. Colour-coding indicates vigilance states. Dark phase is indicated with a dark bar above the graphs. **c)** Representative EEG and EMG traces across vigilance states from individual WT and *Vamp2*^*rlss*^ mice. **d)** Time spent in each vigilance state over 24h for WT (grey) and *Vamp2*^*rlss*^ (red) mice. **e)** Percentage of total sleep time spent in REM sleep. **f)** Number of wake-to-NREM transitions across 24h. **g)** Mean duration of wake episodes for light and dark phases. **h)** Number of NREM-to-REM transitions over 24h. **i)** Percentage of NREM-to-REM transitions (normalised against total number of NREM episodes) over 24h. **j)** Number of wake-to-NREM and NREM-to-REM transitions over 24-h. For figures **f-j**, the minimum length of NREM and wake episodes considered is 1 min, with no minimum duration for REM episodes. **k)** Distribution of NREM episode durations over 24h. Here, all NREM episodes longer than 16 seconds are included. **I)** Distribution of REM episode durations over 24h. Individual data points are shown as is mean ± SEM, p<0.05*, p<0.01**, p<0.01***.

Investigations into the nature of the *Vamp2*^*rlss*^ mutation were driven by observations that this allele was phenotypically distinct from either heterozygous or homozygous null mutant mice (*11*). Western blots from whole brain, cortex or hippocampus indicated that the mutant protein was not as stable as wildtype, with levels ranging from 25-65% of controls. Notably the associated tSNARE protein syntaxin 1a (STX1A) was unaffected (**Fig. 3A, Fig S3A,B**). Immunofluorescence labelling of hippocampal neuronal cultures confirmed that synaptic VAMP2 levels, when normalised to that of the synaptic active zone marker BASSOON, were about 50% that of controls (**Fig. 3 B,C, Fig. S4 C-E**) and these deficits were mirrored in synaptic fractions from cortex or hippocampus (**Fig. 3 C,D**). We investigated whether residual mutant VAMP2 protein was functionally anomalous. Based on evidence that mutations in the transmembrane domain of VAMP2 may affect vesicle fusion, fusion pore dynamics and exocytosis (*12-14*), we conducted reciprocal immunoprecipitations using antibodies to VAMP2 or STX1A. Using either native protein or tagged expression protein for immunoprecipitation, we were unable to detect a consistent difference in protein interaction (**Fig. S3E-H**) although neither may reflect the true dynamism of the interaction between vSNARE and tSNARE components in the context of their physiological lipid environment. Elsewhere, we detected what may be a compensatory homeostatic response in mutants. In cultured neurons, dendritic branching and synaptic density were higher in mutants (**Fig. S4A,B**). Golgi-stained brain sections also revealed an increase in dendritic spine count (**Fig. 3D,E**) with greater numbers of immature spines in mutants (**Fig S5**). At an ultrastructural level, no gross differences in hippocampal synaptic measures were identified. However, the density of synaptic vesicles and the density of docked vesicles were ∼2-fold greater in homozygotes (**Fig. 3F-H**).

**Figure 3.**
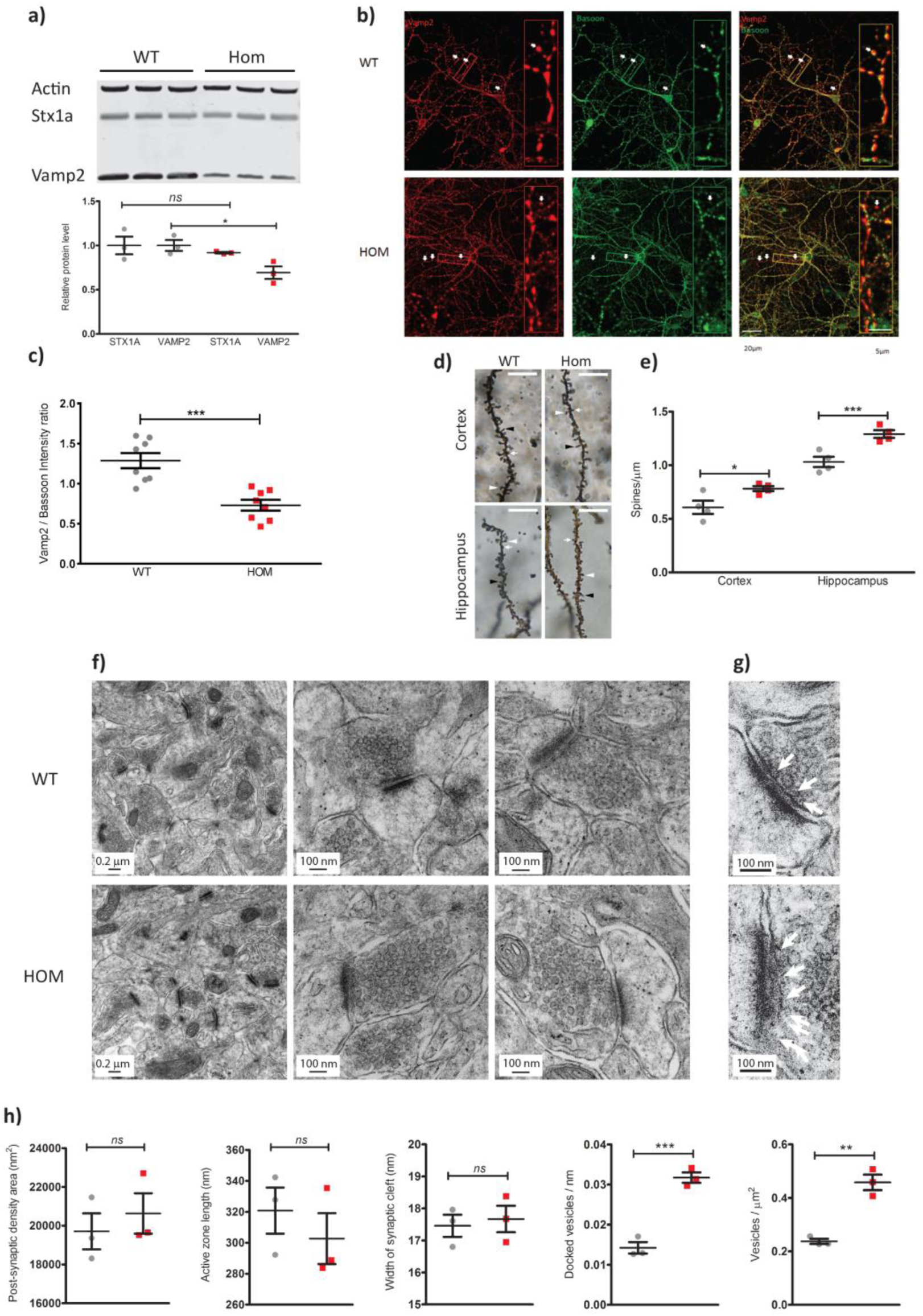
Molecular and cellular deficits in *Vamp2*^*rlss*^. **a)** Western blots of whole brain lysates in wildtypes (WT, grey) and homozygotes (*Vamp2*^*rlss*^, red). STX1a and VAMP2 are quantitated relative to actin levels. **b)** WT and *Vamp2*^*rlss*^ hippocampal cell cultures immunostained with VAMP2 (red), BASSOON (green), co-localization shown in merged channel images (yellow, right panels). Arrows indicate pre-synaptic active zone containing both VAMP2 and BASSOON. Insert shows a close up of stained neuron showing VAMP2, BASSOON and co-localization. Scale Bars as shown. **c)** VAMP2 expression in DIV15 primary cell culture. VAMP2 fluorescence intensity was normalized to that of BASSOON. **d)** Golgi-stained sections showing mushroom (white arrowhead), long (black arrowhead) and stubby (white arrow) spines. Scale bar, 10 µm. e) Spine counts per unit length in WT (grey) and *Vamp2*^*rlss*^ (red) samples. **f)** Electron micrographs of hippocampal sections. Scale bars as shown. **g)** Close-up of images in **h)** showing docked vesicles (arrows). Scale bars as shown. j) Measurement of synaptic parameters. Individual data points are shown as is mean ± SEM, p<0.05*, p<0.01**, p<0.01***.

We tested the effect of *Vamp2*^*rlss*^ on synaptic vesicle release and functional vesicular pool sizes in hippocampal synapses in neuronal cultures. Cultured neurons were transduced with the fluorescence vesicular release reporter sypHy (*15*). Active synaptic boutons were identified using a short burst of high frequency stimulation (20 APs × 100 Hz) which triggers exocytosis of the readily releasable pool (RRP) of vesicles. Fluorescence responses to single APs were next measured in identified presynaptic boutons (**Fig. 4A,B**) enabling an estimate of the average release probability (*pv*) of individual RRP vesicles (*16*). Whilst functional RRP size was similar, *pv* in response to a single AP was profoundly decreased in *Vamp2*^*rlss*^ neurons (**Fig. 4C**). Furthermore, in agreement with electron microscopy data we observed a ∼2-fold increase of the total recycling pool size (TRP) and the total number of SV in *Vamp2*^*rlss*^ neurons as measured by fluorescence sypHy responses to high K+ and NH4Cl containing solutions respectively (**Fig. 4D**). Decreases in *pv* are normally associated with an increase in short-term synaptic facilitation (*17*). Indeed, experiments in acute hippocampal slices revealed a ∼2-fold increase in short term facilitation in *Vamp2*^*rlss*^ slices during short high frequency stimulation of the Shaffer collateral synapses (**Fig. 4e,f**).

**Figure 4.**
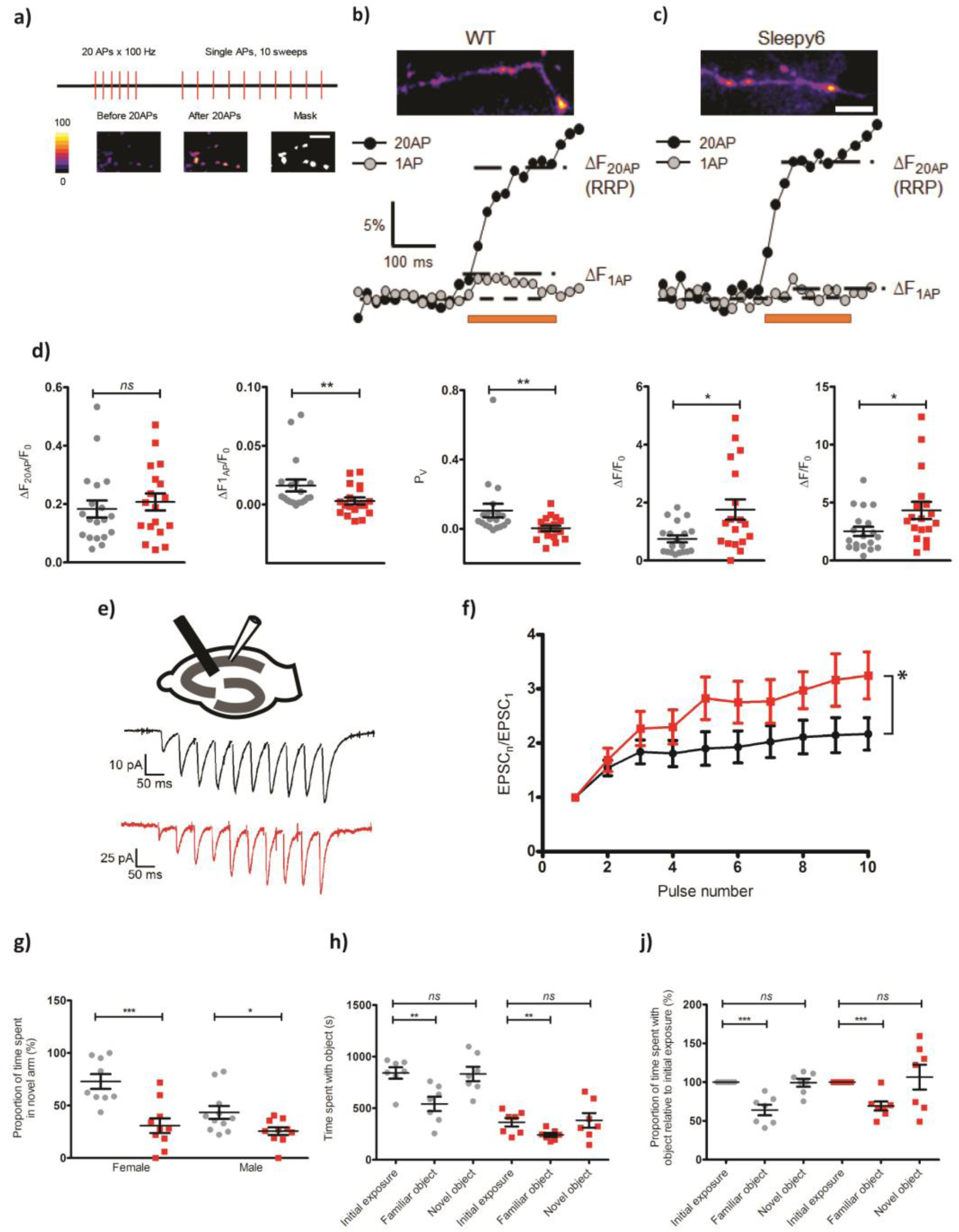
Electrophysiological and behavioural anomalies in *Vamp2*^*rlss*^. **a)** Relative RRP size was determined by measuring the change in fluorescence associated with 20 AP stimuli. This was followed by measuring responses to single AP stimulation. Average traces of sypHy responses from 20-40 boutons to a 20AP × 100Hz burst (ΔF_20AP_ black, beginning of stimulation indicated by arrow) and a single AP (ΔF_1AP_ grey, average of 10 sweeps) in WT **b)** and mutant c) neurons. Average release probability of individual RRP vesicles was estimated in each experiment as *p*_*v*_ = ΔF_1AP_ / ΔF_20AP_. **d)** Comparison of average RRP size, single AP responses, *p*_*v*_, relative total recycling pool (TRP) of vesicles and the total number of SVs in WT (grey) and *Vamp2*^*rlss*^ (red) neurons. Note dramatic reduction in pv in response to a single AP in mutant neurons. n=19 coverslips for WT and n=18 coverslips for Hom from n=5 independent cultures. **e)** Experimental schematics and representative EPSC responses to 10 AP × 20 Hz stimulation in WT (black) and *Vamp2*^*rlss*^ (red) CA1 pyramidal cells. **f)** Comparison of short-term synaptic facilitation. WT n = 8, mutant n = 7. **g)** Percentage time spent in the novel arm of a Y-maze. **h)** Absolute times spent with object in novel object task. **j)** Proportion of time spent with object relative to time spent with original object in novel object task. Individual data points are shown as is mean ± SEM, *p < 0.05, **p < 0.01, ***p < 0.001.

Given the role of VAMP2 in fundamental neural mechanisms and the electrophysiological deficits seen in homozygotes, we expected mutants to display behavioural and sensory discrepancies in addition to changes in sleep. Surprisingly, performance in a wide test battery indicated only mild behavioural anomalies (**Fig. S6**) while sensory function was unchanged or even improved (**Fig. S7,S8**). However, homozygous behaviour in a number of tests pointed to deficits in working memory (Y-maze and Novel Object Recognition), attention (Marble burying, Novel object recognition) and social instability (Social dominance tube test) (**Fig. 4G-J**). These irregularities were defined further through display of numerous stereotypical behaviours analysed using home-cage continuous video monitoring (**Movie S1**).

The sleep deficit identified here in *Vamp2*^*rlss*^ mice uncovers a previously unrecognised role for VAMP2 in sleep and may indicate a critical function for the transmembrane domain in which the mutation occurs. VAMP2 function has been profoundly well-characterised with respect to its fundamental role in vesicle fusion and exocytosis (*18*). Nevertheless, its functional characterisation in mammals *in vivo* has been hampered as the *Vamp2*^*-/-*^ mouse mutant is perinatal lethal while *Vamp2*^*+/-*^ show no behavioural phenotype beyond mild improvements in rotarod performance (*19*). The recent identification of multiple *de novo* variants of *VAMP2* in humans has highlighted how mutations affecting the SNARE zippering mechanism can have severe developmental consequences (*20*). In contrast, the relatively mild consequences of the *Vamp2*^*rlss*^ mutation in mice has enabled us to examine features of VAMP2 function in adults. Moreover, recent publications (*21, 22*) identified *VAMP2* lead SNP associations for chronotype and sleep duration measures suggesting that subtle variations in VAMP2 function could influence sleep quality in humans.

*Vamp2*^*rlss*^ affords a new model to study the genetic mechanisms regulating vigilance state switching. The search for the key “nodes” in the brain which are responsible for instantaneous sleep-wake transitions have undoubtedly provided important insights into these mechanisms (*23-25*). However, as *Vamp2*^*rlss*^ mice display an increased stability of all vigilance states, it is possible that globally impaired vesicular release triggers an inherent inertia in their neuronal circuitry. Thus, it remains a possibility that sleep-wake states are controlled in a global manner, across a distributed circuitry, where the interactions between neurons and regions are tuned by global changes in fundamental neuronal properties such as synaptic vesicular release.

## Methods

### Animals

Animal studies were performed under guidance from the Medical Research Council in Responsibility in the Use of Animals for Medical Research (July 1993), the University of Oxford’s Policy on the Use of Animals in Scientific Research and Home Office Project Licences 30/2686 and 80/2310. When not tested, mice were housed in individually ventilated cages under 12/12 hours light/dark conditions with food and water available *ad libitum*. Inbred strains and mutant colonies were maintained at MRC Harwell and cohorts were shipped as required. ENU mutagenesis and animal breeding regimes were performed as previously reported (*8*). Phenotyping was performed on mouse cohorts that were partially or completely congenic on the C57BL/6J Or C3H.Pde6b+ (*26*) background.

### Video tracking screen for immobility-defined sleep

Video tracking was performed as described (*7*). Briefly, mice were singly housed and placed in light controlled chambers with CCD cameras positioned above the cages (Maplin, UK). Monitoring during the dark was performed using infrared illumination. Analysis of videos by ANYmaze software (Stoelting) was used to track mouse mobility and immobility-defined sleep (successive periods of >40s immobility). Mice for screening were provided by a large-scale ENU mutagenesis project (*8*).

### Gene mapping and sequencing

Affected individuals from G3 pedigrees were used for mapping and mutation detection. Mutations were mapped utilising the Illumina GoldenGate Mouse Medium Density Linkage Panel (Gen-Probe Life Sciences Ltd, UK), providing a map position resolution of about ∼20Mb. Following mapping, whole genome sequencing of the G1 founder male was carried out as described (*8*), variants were confirmed by Sanger sequencing.

### EEG surgery, recording and analysis

Average ages and weights of mice at the time of surgery were 23.8 ± 1.2 weeks & 31.9 ± 0.8 g and 14.7 ± 1.5 weeks & 31.7 ± 1.5 g for *Vamp2*^*rlss*^ and WT mice, respectively. Age differences were permitted so that animals were matched by weight. Previous data indicates that this age difference does not contribute to the differences in sleep observed (*27,28*). *Vamp2*^*rlss*^ mice were fed a high-nutrient diet in addition to regular pellets. Surgical methods, including drug administration and aseptic techniques, were as described (*28*). EEG screws (Fine Science Tools Inc.) were placed in the frontal (anteroposterior +2 mm, mediolateral 2 mm) and occipital (anteroposterior −3.5 to −4 mm, mediolateral 2.5 mm) cortical regions, reference and ground screws were implanted above the cerebellum and contralaterally to the occipital screw, respectively. Two stainless steel wires were inserted in the neck muscle for EMG. Mice were singly housed post-surgery.

After 2 weeks of recovery, mice were placed in cages (20.3 × 32 × 35 cm^3^) in ventilated, sound-attenuated Faraday chambers (Campden Instruments, UK) under a 12h:12h light-dark cycle, with food and water available *ad libitum*. EEG and EMG data were filtered (0.1-100 Hz), amplified (PZ5 NeuroDigitizer preamplifier, Tucker-Davis Technologies) and stored on a local computer (sampling rate: 256.9 Hz). Signals were extracted and converted using custom-written Matlab (The MathWorks Inc., USA) scripts and the open-source Neurotraces software. Vigilance states – wake, NREM, REM, or brief awakenings (short arousals ≤16 s during sleep) – were scored manually using the SleepSign software (Kissei Comtec Co, Nagano, Japan). EEG power density spectra were computed by a Fast Fourier Transform (Hanning window; 0.25-Hz resolution). More detailed recording and initial analysis methods were as published (*28*). Unless otherwise stated, NREM and wake episodes were defined as episodes of at least 1-min duration, allowing brief interruptions ≤16 s (e.g. brief awakenings or brief REM attempts within NREM). REM sleep episodes could be as short as 4 s and bear interruptions ≤4 s.

### Behavioural phenotyping

#### Circadian wheel running

Circadian wheel running was performed as previously reported (*29*). Briefly, mice were singly housed in cages containing running wheels, placed in light controlled chambers and wheel running activity monitored via ClockLab (Actimetrics). Animals were monitored for five days in a 12 hour light/dark cycle followed by twelve days in constant darkness.

#### Passive infrared analysis

Mice were additionally analysed for circadian activity using the COMPASS passive infrared system as described (*9*). Animals were monitored for four days in a 12 hour light/dark cycle followed by ten days in constant darkness.

#### Open field behaviour

Mice were placed into one corner of a walled arena (45cm X 45cm) and allowed to explore on two consecutive days for 30 minutes (*30*). Animal movements and position were tracked using EthoVision XT analysis software (Noldus).

#### Light/Dark box

Individual mice were placed into one corner of an enclosed arena separated into light and dark compartments (*31*). Over 20 minutes animal movements and position were monitored by EthoVision XT.

#### Marble burying

Briefly, a cage was prepared with approximately 5cm deep sawdust bedding (*31*). 9 marbles were placed on the surface of the sawdust, evenly spaced in a regular pattern. The mouse was introduced and left in the cage with the marbles for 30 minutes. After 30 minutes the number of marbles remaining unburied, partially buried or completely buried was counted. Statistical differences were determined using the Mann-Whitney test.

#### Acoustic startle response (ASR) and prepulse inhibition (PPI)

ASR and PPI were measured as in (*32*). Mice were placed in the apparatus (Med Associates, VT, USA) and responses to sound stimuli were measured via accelerometer.

#### Novel object recognition

A modified version of the novel object recognition task adopted for use in the home cage was used using video analysis of behaviours. On day one, animals were presented with one novel object (glass jar or lego bars) in a corner of their home-cage for 30 minutes at the beginning of the dark phase. On the following night the same object was introduced at the same position in the home-cage, again at the beginning of the dark phase for 30 min. On the third night a second object, different to the first one, was introduced at a different position in the home-cage but at the same time of night and for the same duration. Time spent inspecting objects was measured manually from video-recordings using a stop watch.

#### Social dominance test

Dominant and submissive behaviours were assessed for pairs of male mice using a specialised Plexiglass tube (*31*). Pairs of mice, one wildtype and one homozygous mutant from different group-housed cages, were placed in the tube and behaviour registered as dominant (“win”) or subordinate. 5 pairs were used and total numbers of “wins” per genotype was recorded. Statistical differences were ascertained using a Chi-Squared test.

#### Y-maze

A forced alternation y-maze test was used to evaluate short term working memory in mice (*31*). Mice were video tracked at all times using Ethovision XT, preference for the novel arm is indicated by an occupancy of greater than 33%.

#### Mechanical sensitivity, von Frey Test

Mechanical sensitivity was assessed as in (*34*) by determining the withdrawal threshold of the hind limb to a mechanical stimulation applied under the foot pad using an electronic von Frey apparatus (MouseMet, TopCat Metrology, UK). 5 consecutive measurements per hind paw were taken and the average of the last 4 were calculated.

#### Heat sensitivity, Hot Plate Test

A hot plate (BioSeb, Chaville, France) set at 51°C was used for this test (*31*). Animals were placed on the plate and the latency to the first paw reflex (withdrawal reflex of one paw) was measured.

#### Optokinetic response

Visual acuity was assessed by head tracking response to a virtual-reality optokinetic system (Cerebral Mechanics Inc) (*33*).

#### Auditory brainstem response (ABR)

Auditory Brainstem Response tests were performed using a click stimulus in addition to frequency-specific tone-burst stimuli as described (*34*). ABRs were collected, amplified and averaged using TDT System 3 (Tucker Davies Technology) driven by BioSig RZ (v5.7.1) software. All stimuli were presented free-field to the right ear of the anaesthetised mouse, starting at 70 dB SPL and decreasing in 5 dB steps. Auditory thresholds were defined as the lowest dB SPL that produced a reproducible ABR trace pattern and were determined visually.

#### Grip strength

Grip strength was assessed using a Grip Strength Meter (BioSeb, Chaville, France). Readings were taken from all four paws, three times per mouse as per manufacturer’s instructions (*30*). Measures were averaged and normalised to body weight.

#### HCA home cage analysis

Group housed animals were monitored as described (*35*). Briefly, group housed mice were tagged with RFID micochips at 9 weeks of age and placed in the Home Cage Analysis system (Actual Analytics, Edinburgh) which captured mouse behaviour using both video tracking and location-tracking using RFID co-ordinates.

### Golgi-Cox Staining, Spine Count Analysis

Brains dissected from 16-week old females were used for analysis. Golgi-Cox neuronal staining was performed using the FD Rapid GolgiStain Kit (FD NeuroTechnologies Inc, USA) according to the manufacturer’s instructions. 100μm sections were taken using a vibratome, mounted upon charged slides, cleared in Histo-Clear (National Diagnostics, UK) and coverslipped. Neurons were viewed on an Axio-Observer Z1 (Zeiss) microscope. Z stack images were processed using extended depth of focus and Zen software (Zeiss). Visualisation and measurements were taken using ImageJ (http://rsbweb.nih.gov/ij/). The number and type of spines on each neurite - stubby, long, mushroom and branched (*36*) - were counted. At least 50 neurites per region per animal were analysed.

### Electron Microscopy

Brains were fixed by cardiac perfusion (buffer solution: sodium cacodylate buffer pH 7.2 containing 4.35% sucrose, fixation solution: 2.5% glutaraldehyde and 4% paraformaldehyde in buffer solution) removed and coronally cut into 350 µm thick sections using a Vibratome (Leica VT1000 S, Leica Biosystems, Nußloch, Germany). Selected regions from hippocampus were dissected, further fixed with osmium tetroxide and uranyl acetate and dehydrated. After embedding in epoxy resin the tissue was cut into 70 nm thick sections and post-stained with uranyl acetate and lead citrate for imaging using a transmission electron microscope FEI Tecnai 12 TEM (FEI, Hillsboro, Oregon, USA). To avoid any bias between wildtype and homozygous synapses, overview pictures were taken and the number of visible synapses in this plane counted by hand. Higher magnification pictures were exported into ImageJ and measurements taken for each synapse. At least 60 synapses were analysed per animal.

### Plasmids

Plasmid constructs were generated using Invitrogen Gateway® Gene Cloning technology. Wild-type (WT) STX1a and VAMP2 coding sequences were first amplified from cDNA libraries with proofreading polymerase (Platinum SuperFi DNA Polymerase, Invitrogen). Mutant VAMP2 (mVAMP2) was amplified using cDNA obtained from mouse brain. Primers for cloning were STX1a: Forward CACCAAGGACCGAACCCAGGA; Reverse CTATCCAAAGATGCCCCCG VAMP2: Forward CACCTCGGCTACCGCTGCCA; Reverse TTAAGTGCTGAAGTAAACGATGA. Amplified DNA fragments were cloned into pENTR™/D-TOPO™ vector. STX1a was subsequently transferred to pcDNA™ 3.1/nV5-DEST Mammalian Expression Vector (V5 Epitope, Invitrogen), and both WT and mutant VAMP2 sequences were shuttled into Gateway™ pDEST™ 26 Vector (6xHis tag, Invitrogen). The in-frame V5-STX1a and 6xHis-(m)VAMP2 sequences were verified by DNA sequencing.

### Antibodies

Primary antibodies for immunoprecipitation and Western blotting included anti-V5 and anti-His (R&D systems), Mouse anti-VAMP2 (SySy Synaptic Systems, Göttingen, Germany) and Rabbit anti-syntaxin (Sigma, St. Louis, Missouri, US). Secondary antibodies were (IRDye 800CW Goat anti-Mouse IgG, 1:15000 and IRDye 680RD Goat anti-Rabbit IgG, 1:15000). Primary antibodies for immunofluorescence studies were Rabbit anti-VAMP2, Guinea-pig anti-BASSOON and Mouse anti-MAP2 (Synaptic systems, 188004; Novus, SAP7F407 and Synaptic systems, 188 004, respectively). The corresponding secondary antibodies included: 488 Donkey anti-mouse antibody (Invitrogen), 596 Goat anti-guinea-pig antibody (Invitrogen), and 647 Goat anti-rabbit antibody (Life Technologies).

### Cell culture and transfection

Human embryonic kidney 293T (Hek293T) cells were maintained in T75 flasks containing Dulbecco’s Modified Eagle Medium (DMEM, Gibco) supplemented with 10% fetal bovine serum (FBS, Gibco) and Penicillin-Streptomycin (Gibco). Prior to transfection, cells were seeded at 2x 10^5^ cells per well in a 6-well plate and left to adhere overnight. Plasmids (total of 2.5 µg) were transfected with Lipofectamine 2000 (Invitrogen) at a ratio of 1:2 in Opti-MEM I Reduced Serum Medium (Gibco™). Co-transfections of V5-STX1a and 6xHis-mVAMP2 were adjusted so that equivalent amounts of the two proteins were expressed in culture. Transfected cells were then incubated at 37°C for 16-24 hours to express tagged proteins.

### Cell and tissue lysis and fractionation

Cells were harvested 24 hours after transfection and lysed by suspension in ice-cold RIPA buffer (Sigma-Aldrich: R0278-50ML) supplemented with protease inhibitor mixture (Roche Diagnostics). Suspensions were incubated at 4°C for 30 mins and centrifuged at 4°C, 14000 rpm for 20 mins. Tissue lysis and crude synaptic fractionation was performed using Syn-PER Synaptic Protein Extraction Reagent (Thermo Scientific) following the manufacturer’s instructions.

### Immunoprecipitation

For immunoprecipitation (IP) of tagged proteins, the Thermo Scientific Pierce™ Crosslink IP kit (Thermo Fisher Scientific: 26147) was used as per manufacturer’s instructions. Briefly, antibodies were cross-linked to Pierce Protein A/G Plus Agarose to avoid antibody contamination at later steps. Total cell extracts containing V5-STX1a and 6xHis-(m)VAMP2 were pre-cleared with Pierce Control Agarose resin in Pierce Spin Columns for 30 mins at 4 °C and then spun to collect the flow-through, discarding the Control Agarose columns. Pre-cleared samples were then added to antibody cross-linked columns and incubated at 4 °C overnight. After incubation, columns were spun and the flow-through was discarded. The columns were then washed with TBST (25 mM Tris-HCl [pH7.4], 138 mM NaCl, 0.05% Tween 20, National Diagnostics: EC-882) and conditioning buffer to remove non-bound components. The tagged protein complexes were then recovered with the elution buffer supplied. Immunoprecipitation of native protein complexes from hippocampal lysates was carried out as described previously (*32*).

### Electrophoresis and Western Blotting

Samples were prepared in NuPAGE LDS Sample Buffer (Invitrogen: NP0007) supplemented with NuPAGE Sample Reducing Agent (Invitrogen™: NP0004) and heated at 70 °C for 10 mins. Equal amounts of protein were loaded and resolved on NuPAGE Bis-Tris 4-12 % gel (Invitrogen: NP032A) and then transferred to nitrocellulose membarane contained in iBlot Gel Transfer Stacks (Invitrogen: NM040319-01) using an iBlot™ Gel Horizontal Transfer Device (Invitrogen: IB21001). After blotting, membranes were incubated with primary antibodies, diluted in TBST containing 5% skimmed milk, at 4 °C overnight. Subsequently, membranes were washed with TBST and incubated with IR700 and IR800 secondary antibodies (LI-COR Biosciences; Lincoln, NE) for 2 hours at room temperature. After further washes, immunoreactive bands were visualized using an Odyssey Infrared Imaging System (LI-COR Biosciences; Lincoln, NE).

### Hippocampal culture and transduction

Hippocampal neuronal cultures were prepared from P0–P1 littermate WT or *Vamp2*^*rlss*^ homozygous pups (*37*), Neurons were plated on poly-L-lysine (1 mg/ml; Sigma-Aldrich) treated 19-mm glass coverslips at a density of 80,000–120,000 per coverslip. Immunofluorescence and sypHy imaging experiments were performed between 15 and 23 days in vitro (DIV). At DIV 7 neurons were transduced at pFU_sypHy lentivirus containing syntophysin-pHluorin cDNA under control of a ubiquitin promoter. The pFU_sypHy plasmid was kindly provided by Dr. A. Maximov (The Scripps Research Institute, La Jolla, USA).

### Immunofluorescence labelling

To evaluate VAMP2 expression and localization, primary hippocampal neurons were fixed and stained after DIV 15 to 23. Hippocampal cultures were first washed in PBS then fixed with 4% PFA for 10min and blocked in a 20% donkey serum blocking solution in TBST for 1 h at room temperature. All primary antibody incubations were conducted over 2 nights at 4°C. Secondary antibody incubations were for 2h at room temperature. All antibodies were applied in 1% blocking serum to minimize antibodies binding to the plastic-ware. Samples were washed 3x with TBST for 10 min at room temperature between steps. After antibody treatment and final washes, coverslips with cells were mounted onto glass slides with mounting medium (ProLong ™, Gold antifade reagent, Invitrogen) then dried in the dark, overnight at 4°C before imaging.

### Image and Sholl analysis

Fluorescence images were acquired at room temperature with an inverted confocal microscope (Zeiss, LMS 710) using a Plan-Apochromat 40x/1.4 (Oil, DIC, M27) objective or 20x/0.8 (M27) objective to get 4 layer z-stacks. For the synaptic intensity experiment, the same settings for laser power, PMT gain and offset and z-stack thickness were used. The pinhole size was set to 1 Airy unit for the shortest wavelength channel and the faintest image. The z-stacks acquired were compressed into single layer images by maximum projection. For intensity quantification, multi-channel fluorescence images were first converted into monochromatic and colour inverted pictures by imageJ. All fluorescence intensity analysis was conducted at the same setting.

For synaptic distribution analysis, conditions were optimized independently in order to capture data from all neurites for a single cell. PMT gain was optimized individually, using 10% area oversaturation on the shortest wavelength as a reference for all channels (n > 12 for each genotype). Then z-stacks acquired were compressed into single layer images by maximum projection. To quantify synaptic protein distribution further, we investigated VAMP2 distribution pattern by analysing VAMP2 containing synaptic projections and associated neuritic branching using a MATLAB-based SynD program (*38*). Multi-channel fluorescence images were acquired using ZEN software and converted into monochromatic and color inverted pictures by ImageJ for further analyses. Statistical comparisons were performed using a Mann-Whitney test with genotype and intensity as independent factors.

### SypHy fluorescence imaging experiments in neuronal cultures

Primary hippocampal neurons were maintained in modified Tyrode solution containing (mM) 125 NaCl, 2.5 KCl, 2 MgCl_2_, 2 CaCl_2_, 30 glucose and 25 HEPES (pH 7.4) supplemented with NBQX (10 µM, Ascent Scientific) and DL-AP5 (50 µM). APs were evoked by field stimulation via platinum bath electrodes separated by 1 cm (12.5–15 V, 1-ms pulses). To estimate the relative TRP size and total numbers of SVs neurons were perfused with Tyrode modified solution containing either 45 mM KCl or 50 mM NH_4_CL respectively. Images were acquired via 63x objective using a Prime 95B CMOS camera (Photometrics) mounted on an inverted Ziess Axiovert 200 microscope equipped with a 488nm excitation LED light source and a 510 long-pass emission filter. Exposure time was 25ms.

### Image and data analysis of sypHy experiments

Images were analysed in ImageJ and MATLAB using custom-written plugins. A binary mask was placed on all varicosities that were stably in focus throughout all trials and responded to 20 AP 100 Hz burst stimulation. To estimate sypHy fluorescence changes induced by 20 AP × 100 Hz, 1AP, KCl and NH_4_CL the difference between the mean of 8 frames before and 8 frames after the stimulus was calculated for all synapses in the field of view (10-100 range). After subtracting the background, the data were normalised to the resting sypHy signal (F_0_).

### Acute slice preparation and electrophysiological recordings

Whole cell electrophysiological recordings of EPSCs in Shaffer collaterals in acute hippocampal slices from 1 – 2 month old mice have been performed as previously described (*39*). The extracellular perfusion solution contained (in mM): NaCl, 119; KCl, 2.5; CaCl_2_, 2.5; MgSO_4_, 1.3; NaH_2_PO_4_, 1.25; NaHCO_3_, 25; glucose, 10. NMDA and GABA_A_ receptors were blocked routinely with 50 µM D-aminophosphonovalerate (D-APV) and 100 µM picrotoxin. A concentric bipolar stimulating electrode (FHC), connected to a constant current stimulator, was placed in the stratum radiatum and 10 pulses where elicited at 20 Hz. Stimulation intensity ranged from 20 – 320 µA and was adjusted to obtain the first EPSC amplitude in the range of 25 – 100 pA. The elicited EPSCs were recorded from pyramidal neurons of CA1 that were whole-cell patch-clamped using 4–6 mΩ resistance recording pipettes and held at −70 mV. The pipette solution contained (in mM): Cs-Gluconate, 125; HEPES, 10; Na-Phosphocreatine, 10; NaCl, 8; Mg-ATP, 4; Na_3_-GTP, 0.3; EGTA, 0.2; TEA-Cl, 5; biocytin, 0.5. Data were acquired using a PCI-6221 interface (National Instruments) and custom software (LabVIEW). Currents were low-pass filtered (4 – 5 kHz) and digitized at 10 – 20 kHz, before analysis.

### Data analysis and statistics

Unless otherwise stated statistical differences were established using a student’s t-test. Behavioural phenotyping, synaptic spine analysis, electron microscopy, western blot quantification and immunohistochemical quantification were analysed using GraphPad Prism 7 (GraphPad Software). EEG sleep analysis was performed using SPSS (IBM Corp). SypHy fluorescence imaging experiments were analysed using GraphPad Prism 7 and SigmaPlot (Systat Software Inc). The electrophysiological recordings were analysed using LabVIEW (National Instruments), PClamp 10 (Molecular Devices) and Matlab (MathWorks) software. Significance level for all analysis was set at P<0.05.

## Acknowledgements

P.M.N. and M.R.B. were supported by the MRC (grant codes MC_U142684173 and MC_UP_1503/2). M.C.C.G. was supported by a BBSRC DTP grant (BB/J014427/1). V.V.V. was funded by a John Fell OUP Research Fund Grant (131/032) and a Wellcome Trust Strategic Award (098461/Z/12/Z). K.V. was supported by the Wellcome Trust (104033/z/14/z) and ERUK project grant (P1806). S.N.P was funded by the BBSRC (BB/I021086/1) and a Wellcome Trust Strategic Award (098461/Z/12/Z).We thank the biomedical and research support staff at the Mammalian Genetics Unit and Mary Lyon Centre, MRC Harwell Institute. We thank S.D.M. Brown for helpful discussion and comments on the manuscript.

## Author contributions

G.T.B., S.N.P. and P.M.N. were responsible for experimental design and strategy. G.T.B. and I.H. conducted initial forward genetics screens. G.T.B., I.H., N.B., C.A., M.B., and R.S.B. conducted additional behavioural screens and assays. I.H., P.L., and G.T.B. conducted and analysed cellular, EM and morphological assays. M.C.C.G., L.A.B., S.H., V.V.V. and S.N.P conducted secondary sleep screens, EEG recording and analysis. M.Y., C.E. and G.T.B. conducted and analysed all protein work. E.T., E.N. and K.V. conducted and analysed all in vitro electrophysiological and imaging experiments. S.W. directed all animal work and logistics. P.M.N. and G.T.B. assembled all data and wrote the manuscript. All authors contributed to writing the manuscript.

---

**Fig. S1.**
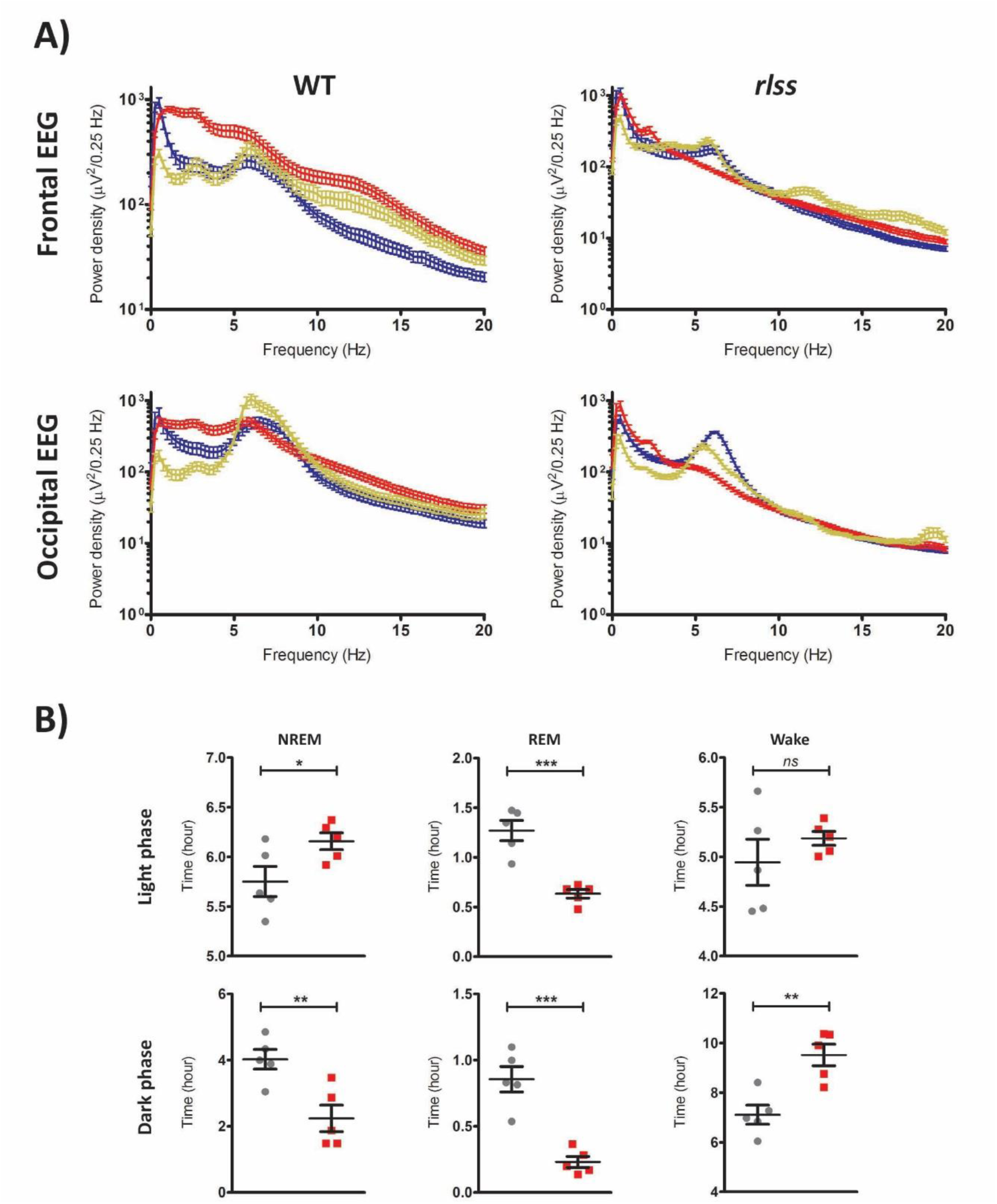
**A)** Mean EEG power density spectra of the different vigilance states - wake (blue), NREM (red) and REM (yellow) - in wild-type (WT) and *rlss* homozygous (HOM) mice calculated from a 24-h baseline recording in the frontal (upper panels) and occipital (lower panels) derivations. **B)** Amount of time spent in each vigilance state during the light and dark phases of a baseline day in wild-type (WT) and *rlss* homozygotes (HOM). n(WT)=5, n(HOM)=5, mean ± SEM, *p < 0.05, **p < 0.01, ***p < 0.001.

**Fig. S2.**
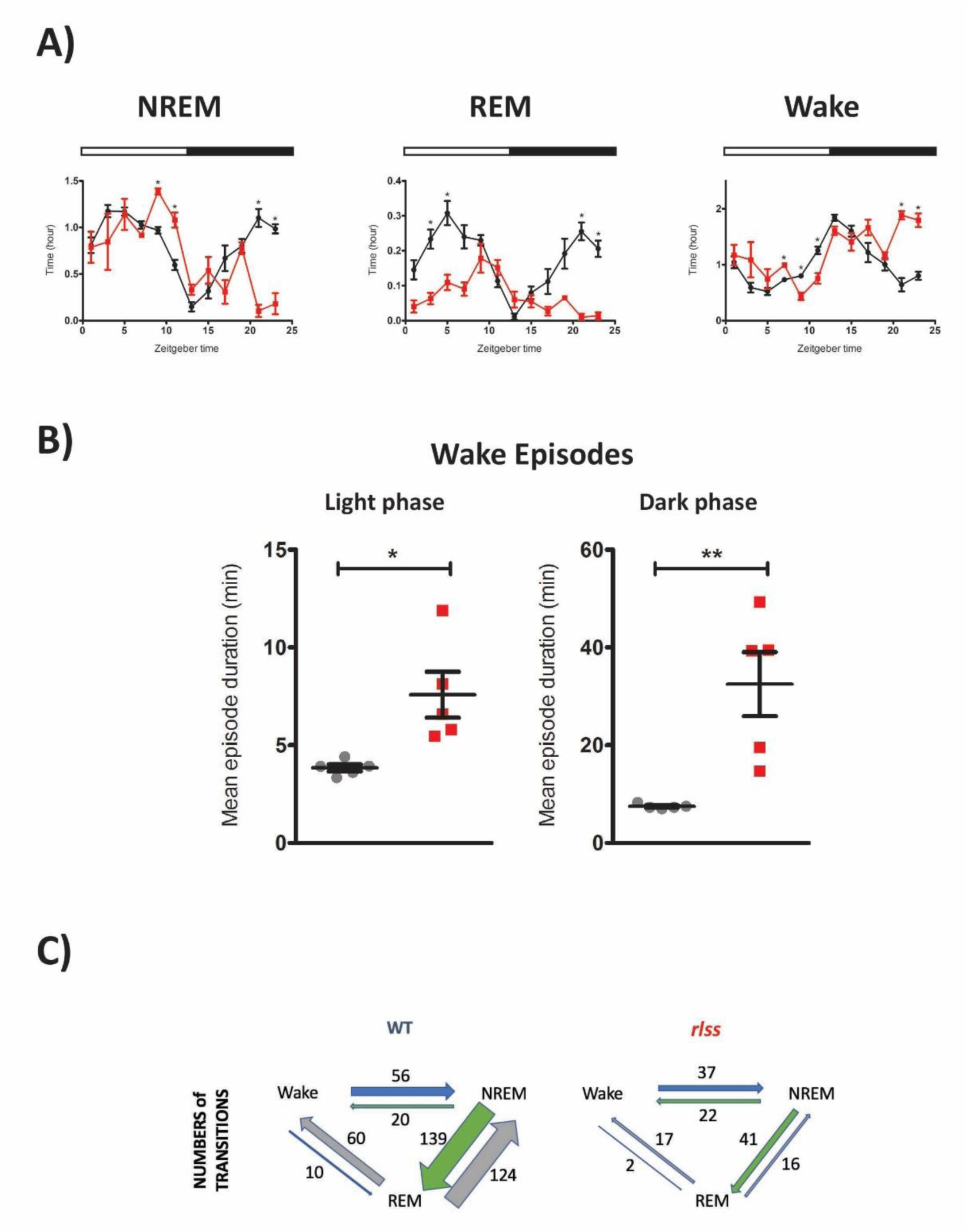
**A)** Distribution of vigilance states across a 24-h baseline day in wild-type (WT) and *rlss* homozygous (HOM) mice. Light and dark phases are indicated above each graph with white and black bars, respectively. Data are plotted in 2-h bins. **B)** Mean duration of wake episodes of at least 16 seconds (as opposed to 1 min in **Fig 2h**) during the 12-h light and dark phases of a baseline day. n(WT)=5, n(HOM)=5. Mean ± SEM. *p < 0.05, **p < 0.01 **C)** Number of vigilance state transitions in WT and *rlss* homozygous (HOM) mice. All NREM and wake episodes of at least 1 min within a 24-h baseline day were included (no minimum duration for REM episodes).

**Fig. S3.**
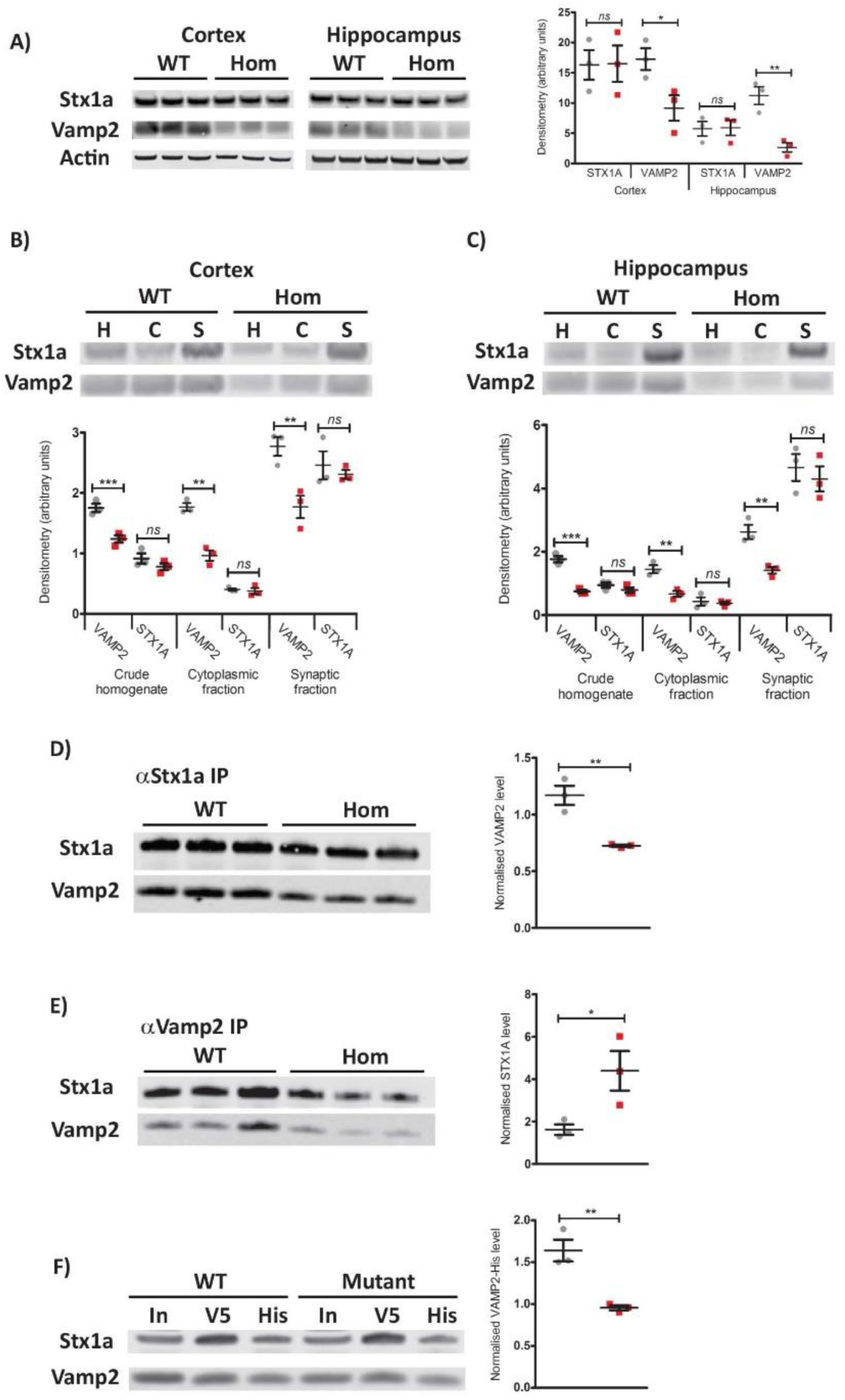
**A)** Western blots (left) and densitometry (right) of cortical and hippocampal whole cell lysates in wildtype (WT, grey) and *rlss* homozygous (Hom, red) samples. Levels of STX1a and VAMP2 are quantitated relative to actin levels. Western blots and densitometry of cortical **B)** and hippocampal **C)** whole cell (W), cytosolic (C) and synaptic (S) cellular fractions. Levels of STX1a and VAMP2 are quantitated relative to actin levels. Western blot and densitometry of IP from hippocampal tissue using either primary Stx1a **D)** or Vamp2 **E)** antibody. Levels of the co-precipitated protein are quantitated relative to the level of the primary antigen precipitated. **F)** Western blots and densitometry of pull downs using approximately equal levels of transfected 6xHis-VAMP2 (WT or mutant) and V5-STX1A proteins in HEK cells. Western blot shows result of a single set of pull downs showing input (In), precipitate using anti-V5 as the primary antibody (V5) and precipitate using anti-His as the primary antibody (His). Densitometry shows quantitation of the co-precipitate relative to the level of the primary antigen precipitated. Individual data points are shown (n=3) as is mean ± SEM, *p < 0.05, **p < 0.01, ***p < 0.001.

**Fig. S4.**
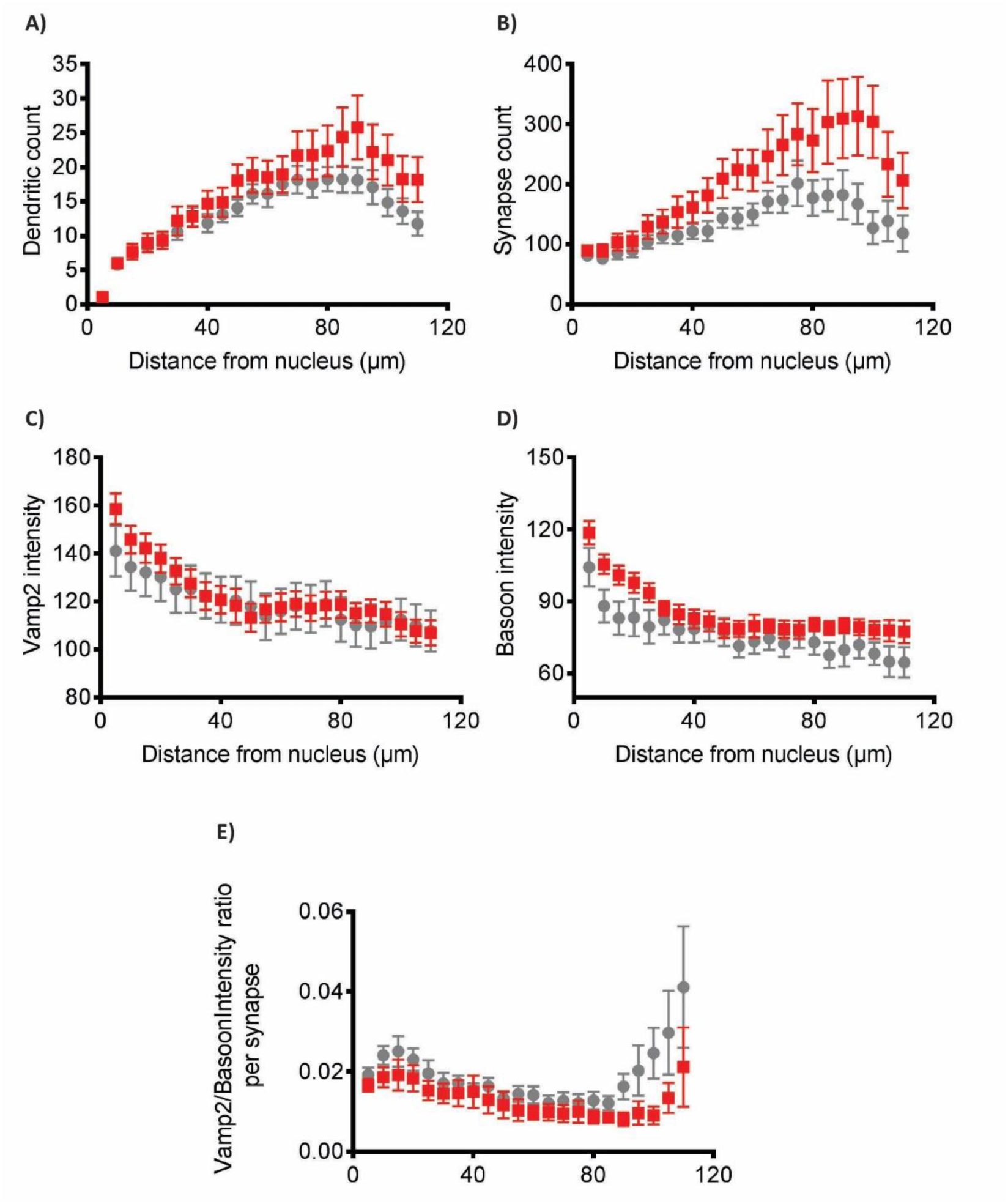
**A)** Dendritic branching in wildtype (grey) and mutant (red) cultures was evaluated by Sholl analysis of Map2 stained hippocampal neurons and dendritic branch counts were plotted against distance from the cell soma base. **B)** The number of pre-synaptic projections were evaluated using Bassoon as a marker and synaptic count plotted against distance from the cell soma base. **C)** Vamp2 immunofluorescence staining intensity plotted against distance from cell soma base. **D)** Bassoon immunofluorescence staining intensity plotted against distance from cell soma base. **E)** Vamp2 fluorescence intensity from **C)** normalized by Bassoon expression in **D)** in each synaptic projection from **B)**. Normalized Vamp2 expression was plotted against distance from soma base. Neurites were detected using SynD software. WT n=14, Hom n = 12. Data are expressed as mean ± SEM.

**Fig. S5.**
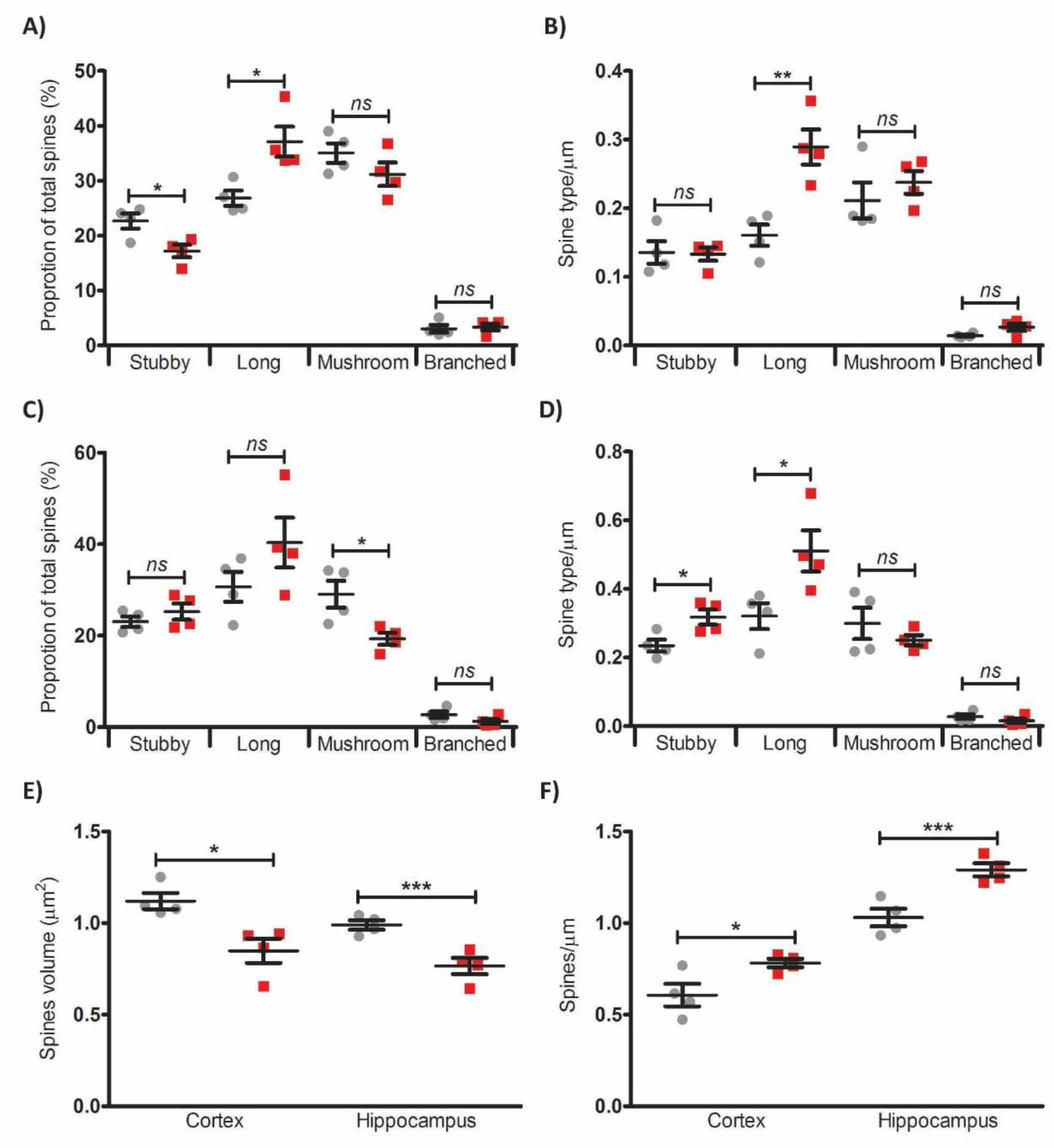
Golgi-Cox analysis of spines. Proportion and number of each spine type respectively in cortical (**A,B**) and hippocampal (**C,D**) sections from wildtype (grey) and *rlss* homozygous (red) mice. Average spine volume (**E**) and spine count per μm (**F**) in cortical and hippocampal sections. Individual data points are shown (n=4) as is mean ± SEM, *p < 0.05, **p < 0.01, ***p < 0.001.

**Fig. S6.**
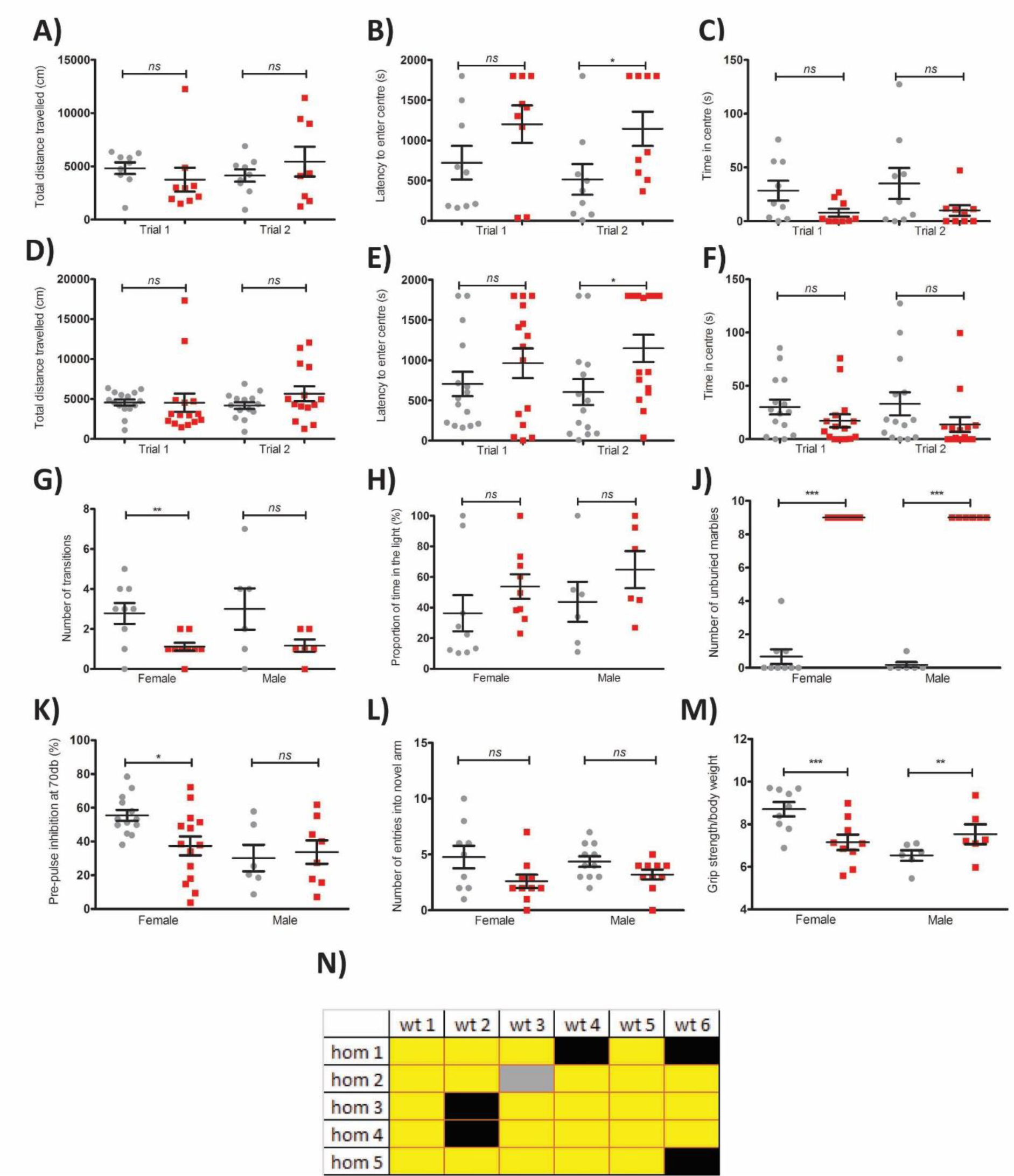
Behavioural test battery measures in wildtype (grey) and *rlss* homozygous (red) mice. Open field distance travelled, latency to centre area and time spent in centre for females (**A,B,C** respectively) and males (**D,E,F** respectively), **G**) number of transitions in a Light-dark box, **H**) proportion of time in dark compartment of a Light-dark box, **J**) numbers of marbles left unburied in marble burying test, **K**) Prepulse inhibition of acoustic startle response, **L**) Y-maze number of entries into novel arm, **M**) Grip strength adjusted to body weight, **N**) Social dominance tube test. Yellow = WT is dominant. Black =inconclusive. Grey = untested. Cohort numbers for individual tests are as shown. Mean ± SEM, **p < 0.01, ***p < 0.001.

**Fig. S7.**
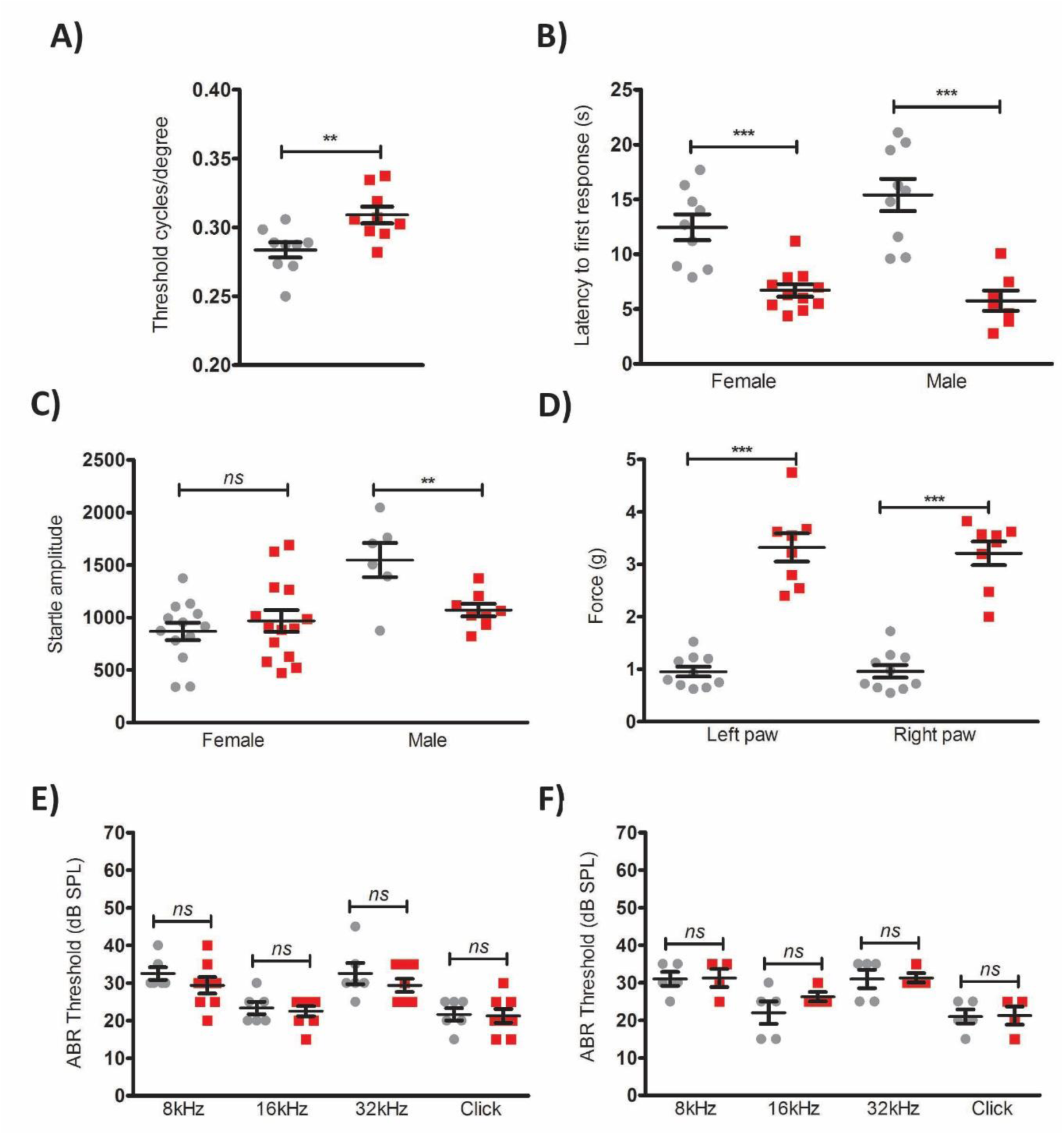
Sensory function in wildtype (grey) and *rlss* homozygous (red) mice. **A**) Visual acuity measured using an optokinetic response drum, **B**) Thermal nociception measuring latency to first response in hot plate test, **C**) Acoustic startle response, **D**) Mechanical sensitivity thresholds using the von Frey test, Acoustic brainstem response (ABR) threshold measures in males **E**) and females **F**). Cohort numbers for individual tests are as shown. Mean ± SEM, **p < 0.01, ***p < 0.001.

**Fig. S8.**
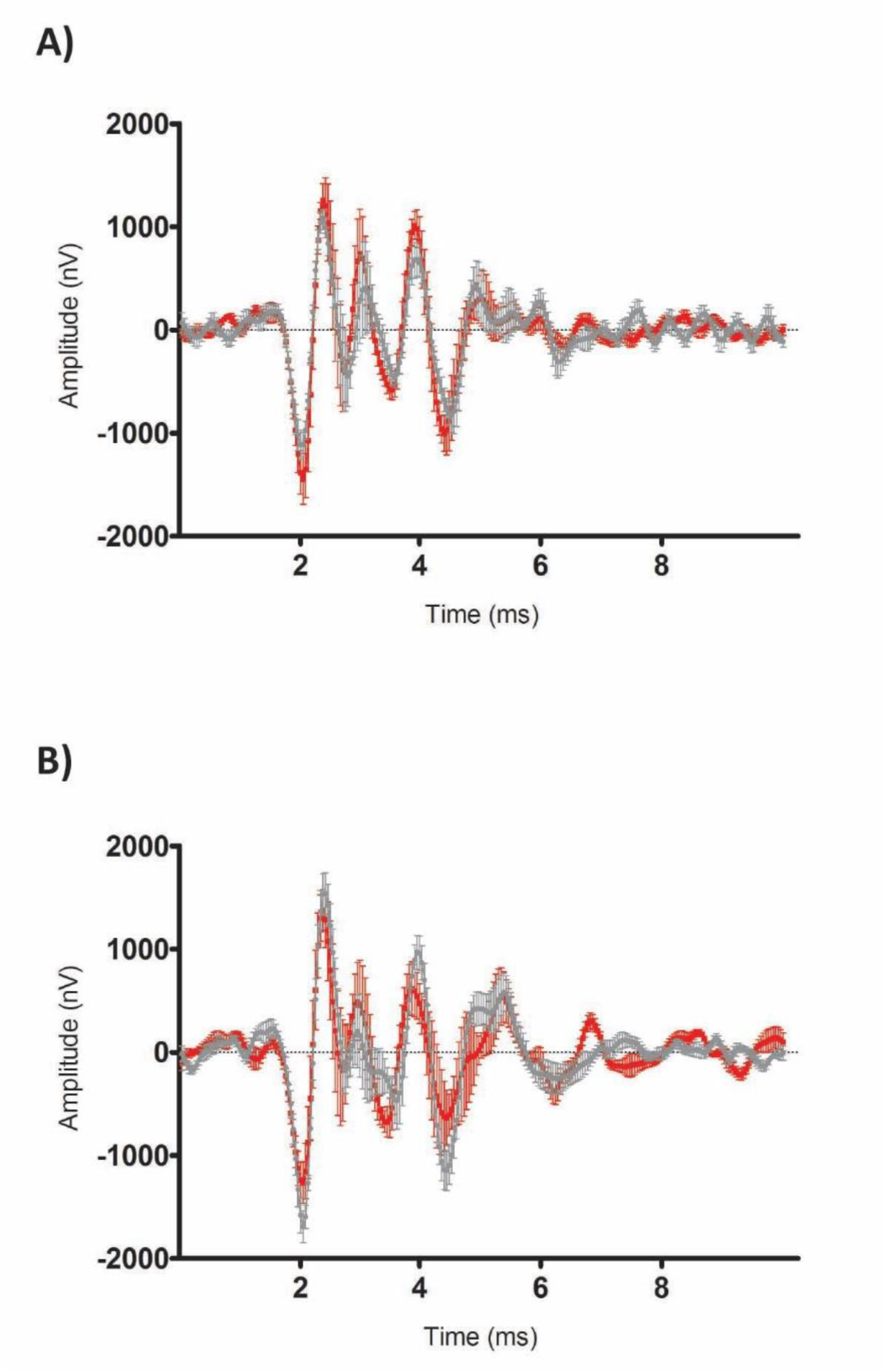
ABR waveform in response to a 70db stimulus in male **(A)** and female **(B)** animals showed no evidence of synaptic malfunction along the auditory pathway. There are no overall differences in the amplitude or latency of ABR components of mutant mice compared to wildtype controls suggesting no likely malfunction of the various peak-generating synapses along the auditory pathway. Traces show mean waveform ± SEM (Male WT, n=5; Male Hom, n=4; Female WT, n=6; Female Hom, n=8).

